# Evaluation of S1PR1/pSTAT3 and S1PR2/FOXP1 Expression in Aggressive, Mature B Cell Lymphomas

**DOI:** 10.1101/472449

**Authors:** Mustafa Al-Kawaaz, Teresa Sanchez, Michael J Kluk

**Affiliations:** Weill Cornell Medicine, Dept. of Pathology and Laboratory Medicine New York, NY, USA

## Abstract

Aggressive, mature B-cell lymphomas represent a heterogeneous group of diseases including Burkitt Lymphoma (BL), High Grade B Cell Lymphomas (HGBL) (eg, Double-Hit B cell lymphomas (HGBL-DH: HGBL with MYC and BCL2 and/or BCL6 translocations)), HGBL, Not Otherwise Specified (HGBL, NOS) and Diffuse Large B Cell Lymphoma. The overlapping morphologic and immunohistochemical features of these lymphomas may pose diagnostic challenges in some cases, and a better understanding of potential diagnostic biomarkers and possible therapeutic targets is needed. Sphingosine 1 Phosphate Receptors (S1PR1-5) represent a family of G-protein coupled receptors that bind the sphingolipid (S1P) and influence migration and survival pathways in a variety of cell types, including lymphocytes. S1PRs are emerging as biomarkers in B cell biology and interaction between S1PR pathways and STAT3 or FOXP1 has been reported, especially in DLBCL. Our aim was to extend the understanding of the S1PR1, STAT3 and S1PR2, FOXP1 expression beyond DLBCL, into additional aggressive, mature B cell lymphomas such as BL, HGBL-DH and HGBL,NOS.

Herein, we report that S1PR1 and S1PR2 showed different patterns of expression in mantle zones and follicle centers in reactive lymphoid tissue and, among the lymphomas in this study, Burkitt lymphomas showed a unique pattern of expression compared to HGBL and DLBCL. Additionally, we found that S1PR1 and S1PR2 expression was typically mutually exclusive and were expressed in a low proportion of cases (predominantly HGBL involving extranodal sites). Lastly, FOXP1 was expressed in a high proportion of the various case types and pSTAT3 was detected in a significant proportion of HGBL and DLBCL cases. Taken together, these findings provide further evidence that S1PR1, pSTAT3, S1PR2 and FOXP1 play a role in a subset of aggressive mature B cell lymphomas.

## Introduction

Aggressive, mature B-cell lymphomas represent a heterogeneous group of diseases including Burkitt Lymphoma (BL), High Grade B Cell Lymphomas (HGBL) (eg, Double-Hit B cell lymphomas (HGBL-DH: HGBL with MYC and BCL2 and/or BCL6 translocations)), HGBL, Not Otherwise Specified (HGBL, NOS)) and Diffuse Large B Cell Lymphoma (DLBCL). DLBCL can be further classified into germinal center B-cell-like (GCB subtype) and activated B-cell-like (ABC; non-GCB subtype)^1, 2^. In some cases, classification of these types of aggressive mature B cell lymphomas can pose diagnostic challenges, and due to their aggressive clinical course, a better understanding of diagnostic biomarkers and potential therapeutic targets is needed.

Sphingosine 1 Phosphate Receptors (S1PR1-5) represent a family of G-protein coupled receptors that bind the sphingolipid (S1P) and influence migration and survival pathways in a variety of cell types; S1PRs are emerging as biomarkers in B cell lymphomas^3–6^ and B cell development^7^. Our prior work, which demonstrated S1PR1 expression in a subset of Classic Hodgkin Lymphoma cases^8^, has recently been supported by others^9^. Additional studies indicate that S1PR expression may influence anatomic location/distribution in a variety of types of lymphoma ^10^. Although the role of S1PR1 and S1PR2 have been examined in mantle cell lymphoma and DLBCL, the role of this S1P pathway in BL and other HGBL, including HGBL-DH has not been specifically characterized.

STAT3 is a transcription factor which regulates tumorigenesis in a variety of lymphoproliferative disorders and is therapeutically targetable^11, 12^. pSTAT3 has been reported to be a potential biomarker in DLBCL which may depend on anatomic location, according to one study^13^. STAT3 was reported to show high levels of expression and phosphorylation in ABC-DLBCL^14^. Interestingly, STAT3 was found to be a direct transcriptional activator of S1PR1 expression in various cell types^15^. In DLBCL, S1PR1 over-expression has been reported in approximately 6-40% of DLBCL and was associated with STAT3 phosphorylation in fresh tissue and as well as a negative prognosis^5, 16^. Additional studies, using primary tumor cells, have also shown phospho-STAT3 activity correlated with increased S1PR1 expression in ABC-DLBCL^17^, and that S1PR1 could activate STAT3 in ABC-DLBCL^17^. Furthermore, inhibition of S1PR1 expression, down-regulated STAT3 activity and caused growth inhibition of lymphoma cells^17^. In BL, phosphorylated STAT3 was reported to be associated with multidrug resistance according to one study^18^, however, the expression of phosphorylated STAT3 does not appear to have been adequately examined using clinical material. In terms of other HGBL, phosphorylated STAT3 has been reported to be associated with MYC and MYC/BCL2 double expression in one study^19^, however, there was no association between phosphorylation of STAT3 with rearrangement of MYC, BCL2 or BCL6.

FOXP1 (Forkhead box protein P1) is a transcription factor that is expressed in a subset of GCB-DLBCL, and to a greater extent in ABC-like DLBCL, and may be associated with a poorer overall survival according to some studies ^13, 20^. According to some studies, FOXP1 expression correlates with MYC expression^20^ and in a limited series of 11 cases of Triple Hit Lymphoma (ie, B Cell Lymphomas with MYC, BCL2 and BCL6 translocations), all 11 cases were positive for FOXP1^21^. However, FOXP1 expression in BL and HGBL-DH has yet to be adequately explored. Furthermore, interestingly, FOXP1 was reported to repress S1PR2 expression in ABC- and GCB-DLBCL cell lines and FOXP1 mRNA expression was inversely correlated with S1PR2 mRNA expression in patient cohorts; furthermore, low S1PR2 mRNA expression, especially together with high FOXP1 mRNA expression, was associated with a negative prognosis^22^. In additional studies, S1PR2 was found to play a role in germinal center confinement of B cells^23^ and mutations in S1PR2 reported in GCB-DLBCLs were found to disrupt the inhibitory functions of this receptor^6, 24, 25^.

Taken together, the prior research suggests a role for S1PR1, pSTAT3, S1PR2 and FOXP1 in distinct subtypes of DLBCL, however, testing for these biomarkers in additional patient cohorts and disease subsets is needed. Therefore, our aim was to extend the understanding of the S1PR1/STAT3 and S1PR2/FOXP1 expression beyond DLBCL into additional aggressive, mature B cell lymphomas such as BL, HGBL-DH and HGBL,NOS.

## Methods

Formalin-fixed paraffin embedded (FFPE) tissue samples from 150 total cases were identified from the Weill Cornell Medicine/New York Presbyterian Hospital, Department of Pathology archives. These cases had been reviewed by subspecialist hematopathologists and the cases were diagnosed using 2008 WHO classification criteria^1^. Immunostaining for S1PR1, pSTAT3, S1PR2 and FOXP1 was performed on the Leica Bond Autostainer (Buffalo Grove, IL.) on paraffin embedded tissue sections. We have previously validated the S1PR1 antibody^8^ (Santa Cruz,SC-25489, H60); the S1PR1 staining conditions in this study were: H2(Tris-EDTA(pH9)) antigen retrieval for 20 minutes; 30 min incubation with primary antibody(dilution of 1:100). S1PR2 (Acris, AP0-1198PU-N) performance was confirmed (Figure 1) and staining conditions were: H2(Tris-EDTA(pH9)) antigen retrieval for 20 minutes; 15 minute incubation with primary antibody (dilution of 1:100). Cases were positive for S1PR1 or S1PR2 when >50% of lymphoma cells showed moderate to strong cytoplasmic or membranous staining. FOXP1 (ABCAM, AB16645) staining conditions were: H1(Citrate(pH6)) antigen retrieval for 30 minutes, 15 minute incubation with primary antibody (dilution 1:400). Cases were positive for FOXP1 when >90% (2/2 level positivity) or 50-90% (1/2 level positivity) of lymphoma cells were positive for nuclear staining. Phospho-STAT3 (Cell Signaling, CST 9145; Tyr705) staining conditions were: H2(Tris-EDTA(pH9)) antigen retrieval for 20 minutes, 60 minute incubation with primary antibody(dilution 1:100). Cases were positive for phospho-STAT3 when >10% of lymphoma cells showed nuclear staining. Phospho-STAT3 showed nuclear positivity in vascular endothelial cells in control tissues as previously reported^5, 16^. Vascular endothelial cell and/or stromal cell staining (internal control) was present for each marker in cases when lymphoma cells were negative. This work was supported by the Translational Research Program at WCMC Pathology and Laboratory Medicine and was approved by the Weill Cornell Institutional Review Board (Protocol #: 1509016528).

**Figure 1:**
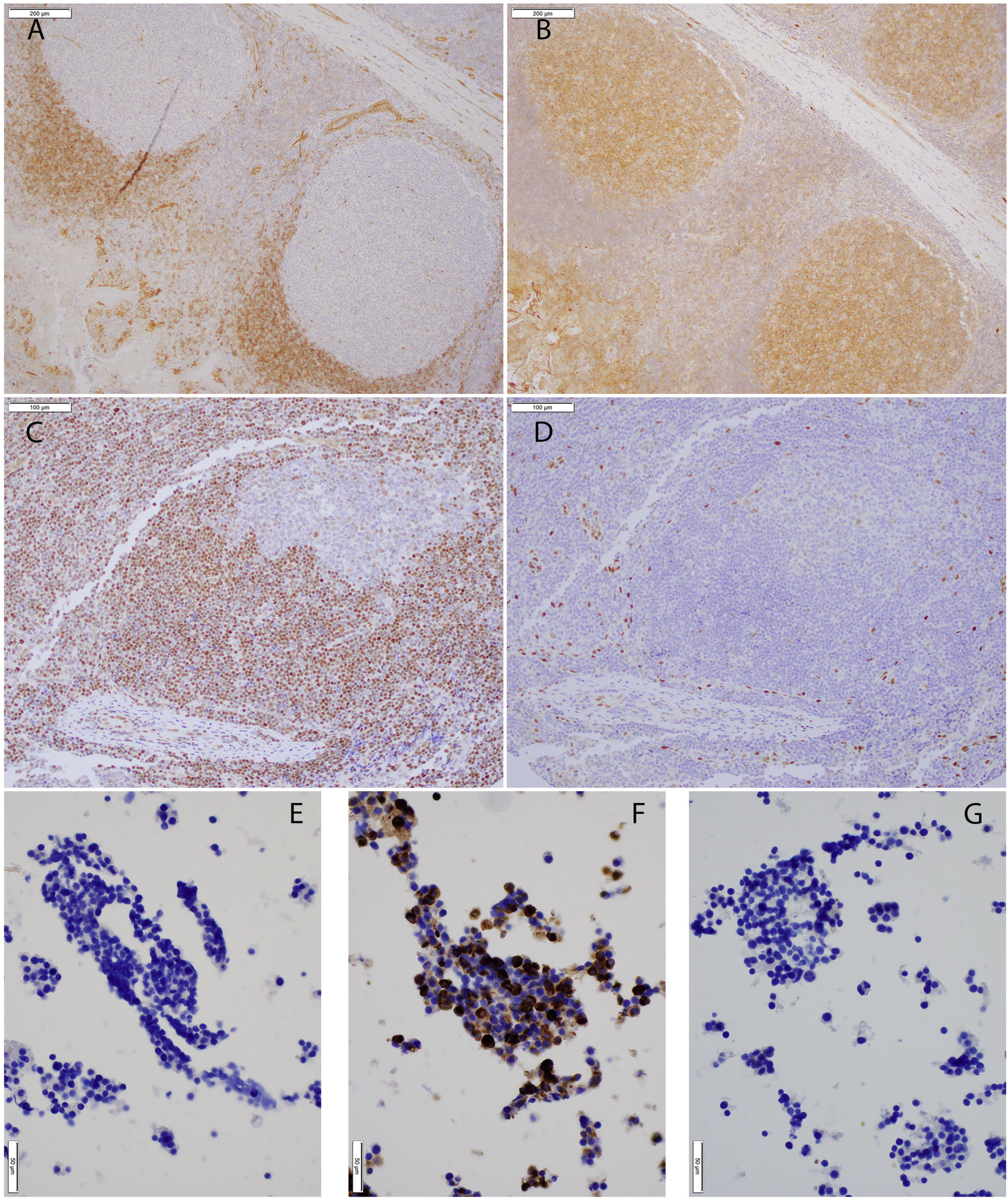
**A-D**: Immunostaining for S1PR1 (**A**), S1PR2 (**B**), FOXP1 (**C**) and pSTAT3 (**D**) in follicles from reactive lymphoid tissue; S1PR1 staining is seen mainly in mantle zones and vascular endothelial cells rather than reactive follicle centers; S1PR2 staining is mainly seen in reactive follicle centers and vascular endothelium, rather than mantle zones; FOXP1 is most highly expressed in mantle zones; pSTAT3 stains scattered cells, including endothelial cells and stromal cells (10x- (**A, B**) or 20x- (**C, D**) magnification; scale bars= 200um (**A, B**) or 100um (**C, D**)). **1, E-G**: Immunostaining for S1PR2 is shown for S1PR1-transfected (**E**), S1PR2-transfected (**F**), and control vector-transfected (**G**) human embryonic kidney 293 cells; S1PR2 staining is appropriately seen for S1PR2-transfected cells, but not for S1PR1- or control vector-transfected cells (40x magnification, scale bars= 50um).

## Results

Immunostaining for S1PR1 showed findings consistent with the reported staining pattern^4^ whereby mantle zones and vascular endothelial cells were S1PR1 positive, while follicle centers were negative for S1PR1 (Figure 1). On the other hand, S1PR2 was expressed more strongly in follicle centers than in mantle zones (Figure 1), which consistent with the S1PR2 mRNA expression studies described previously in mice^23^ and sorted human tonsillar B cells^24^; in addition, S1PR2-staining was specific for S1PR2 when tested with control cell lines (Figure 1). FOXP1 staining was consistent with described patterns^20^ such that FOXP1 was more strongly expressed in mantle zones than follicle centers (Figure 1). Lastly, phospho-STAT3 showed scattered staining in reactive follicles and interfollicular areas, including vascular endothelial cells and scattered stromal cells (Figure 1), consistent with reported staining patterns^5, 16^.

A total of 150 cases of aggressive B cell lymphoma were tested (43 cases of DLBCL, GCB Type; 48 cases of DLBCL, Non-GCB Type; 17 cases of DLBCL, NA (ie, subtyping not available); 11 cases of Double Hit, High Grade B Cell Lymphoma (HGBL-DH); 15 cases of High Grade B Cell Lymphoma, NOS (HGBL, NOS); 15 cases of Burkitt lymphoma (BL), 1 case of Burkitt-like Lymphoma with Del11q) and staining results are summarized in Table 1. BL cases were negative for S1PR1, pSTAT3, and S1PR2, but were uniformly positive for FOXP1 (15/15 cases, 100%). HGBL-DH were also uniformly positive for FOXP1 (11/11 cases, 100%) and mostly negative for S1PR1, pSTAT3 and S1PR2, similar to Burkitt Lymphoma; however, a small subset of the HGBCL-DH cases were positive for pSTAT3 (2/11 cases, 18%), S1PR1 (1/11 cases, 9%) and S1PR2 (1/11 cases, 9%). Interestingly, HGBL, NOS cases also showed near uniform expression of FOXP1 (14/15 cases, 93%), but showed proportionally higher percentages of pSTAT3 (7/15 cases, 47%), S1PR1 (2/15 cases, 13%) and S1PR2 (4/15 cases, 27%) positive cases. Lastly, DLBCL cases were analyzed according to GCB and Non-BCB subtypes according to the Hans algorithm. DLBCL, GCB-type cases showed a significantly lower proportion of FOXP1 positive cases (27/38 cases, 71%) than the HGBL group (25/26, 96%) (Fisher’s exact test, two tailed P value =0.0196). The DLBCL, GCB-Type cases also showed pSTAT3 positivity in 22% of cases (9/40 cases) and were uniformly negative for S1PR1 and S1PR2. The DLBCL, Non-GCB Type cases showed a slightly higher proportion of FOXP1 positive cases (40/47 cases, 85%) compared to DLBCL, GCB Type cases, but this difference was not significant (Fisher’s exact test, two tailed p value=0.1810). The proportion of FOXP1 positive DLBCL, Non-GCB cases was also not significantly different from the HGBL group (Fisher’s exact test, two tailed P value =0.2453). The DLBCL, Non-GCB type cases showed pSTAT3 positivity in 52% of cases (24/46 cases), which was significantly higher than the DLBCL, GCB-Type group (9/40 cases, 22%)(Fisher’s exact test, two tailed P value = 0.0073), and was not significantly different than HGBL group (9/26 cases, 35%)(Fisher’s exact test, two tailed P value = 0.2184). The GCB-type and Non-GCB-type subgroups of DLBCL did not show significant levels of S1PR1 or S1PR2 expression.

**TABLE 1:**
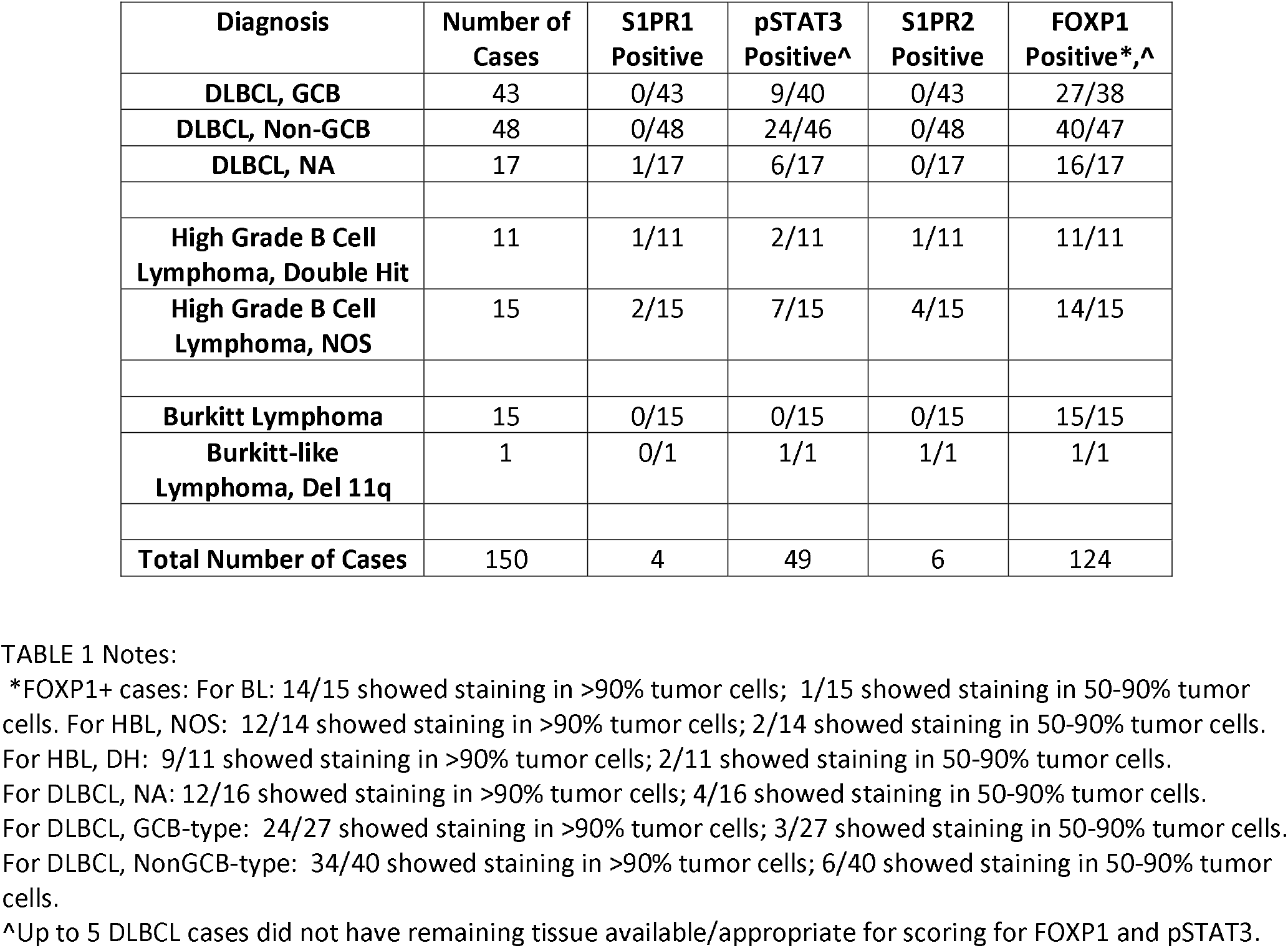
Summary of Immunostaining Results.

Of all cases, 6% (9/150) were positive for S1PR1 and/or S1PR2: 2% (3/150) of cases were S1PR1+, 3% (5/150) were S1PR2+ and 1 case (1/150, <1%) was positive for both S1PR1 and S1PR2. Thus, S1PR1 and S1PR2 staining was mutually exclusive in 89% (8/9) of these cases. The features of the S1PR1+ and S1PR2+ cases are described in Table 2. Such cases were predominantly HGBL. Due to the low prevalence of S1PR1+ and S1PR2+ cases, the respective relationship to pSTAT3 and FOXP1 staining was not possible to ascertain. The four S1PR1+ cases included both pSTAT3+ (2/4) and pSTAT3-negative subgroups (2/4). The six S1PR2+ cases were all FOXP1 positive (6/6).

**TABLE 2:**
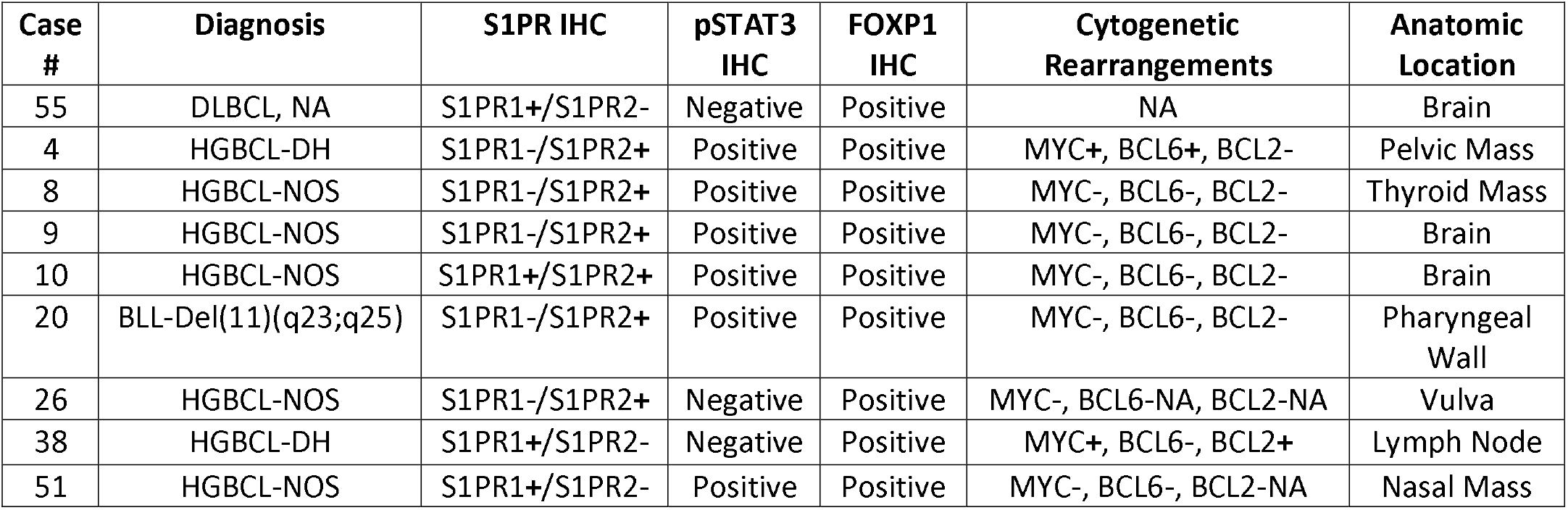
Details of S1PR1+ and S1PR2+ Cases.

| Case # | Diagnosis | S1PR IHC | pSTAT3 IHC | FOXP1 IHC | Cytogenetic Rearrangements | Anatomic Location |
| --- | --- | --- | --- | --- | --- | --- |
| 55 | DLBCL, NA | S1PR1+/S1PR2- | Negative | Positive | NA | Brain |
| 4 | HGBCL-DH | S1PR1-/S1PR2+ | Positive | Positive | MYC+, BCL6+, BCL2- | Pelvic Mass |
| 8 | HGBCL-NOS | S1PR1-/S1PR2+ | Positive | Positive | MYC-, BCL6-, BCL2- | Thyroid Mass |
| 9 | HGBCL-NOS | S1PR1-/S1PR2+ | Positive | Positive | MYC-, BCL6-, BCL2- | Brain |
| 10 | HGBCL-NOS | S1PR1+/S1PR2+ | Positive | Positive | MYC-, BCL6-, BCL2- | Brain |
| 20 | BLL-Del(11)(q23;q25) | S1PR1-/S1PR2+ | Positive | Positive | MYC-, BCL6-, BCL2- | Pharyngeal Wall |
| 26 | HGBCL-NOS | S1PR1-/S1PR2+ | Negative | Positive | MYC-, BCL6-NA, BCL2-NA | Vulva |
| 38 | HGBCL-DH | S1PR1+/S1PR2- | Negative | Positive | MYC+, BCL6-, BCL2+ | Lymph Node |
| 51 | HGBCL-NOS | S1PR1+/S1PR2- | Positive | Positive | MYC-, BCL6-, BCL2-NA | Nasal Mass |

## Discussion

Our study examined the expression of S1PR1, pSTAT3, S1PR2 and FOXP1 using immunostaining of formalin-fixed paraffin embedded patient samples from a clinical cohort of a variety of aggressive B cell lymphoma cases. This was done in an attempt to extend our understanding of these markers beyond DLBCL, given that these potentially important pathways have not been adequately studied in BL and HGBL-DH, etc.

In prior studies of DLBCL, Koresawa et al^5^ reported 13% of their DLBCL cases were S1PR1 positive and were enriched for DLBCL, Non-GCB Type; interestingly they found that primary testicular DLBCL cases showed a higher prevalence of S1PR1 positivity (54%). Thus, the overall prevalence of S1PR1+ cases in their cohort was approximately 10%, when considering only DLBCL cases which were not primary testicular DLBCL. They found that among early stage DLBCL patients, S1PR1 expression was associated with a poor prognosis. Paik et al^16^ reported S1PR1 positivity in 40% of their DLBCL cases and the S1PR1+ cases also appeared enriched for DLBCL occurring in extranodal sites, but there was no significant difference for S1PR expression between the GCB-type and Non-GCB-type subgroups; Paik et al also found that S1PR1 expression was associated with a poor prognosis. Lastly, Nishimura et al^4^ reported that 6% of DLBCL cases were S1PR1 positive. The differences in the prevalence of S1PR1 positivity among the different cohorts reported may have been due to the variability of cytoplasmic/membrane staining patterns observed^5^ as well as differences in the proportion of primary testicular DLBCL cases among the cohorts. In our study, we have found that only approximately 1% (1/108) of DLBCLs were positive for S1PR1. However, we found that S1PR1 was expressed in approximately 11% (3/26) of HGBLs. The prevalence of S1PR1+ lymphomas in these groups in our study is most in line with the prevalence of S1PR1 positivity reported by Koresawa et al^5^ and Nishimura et al^4^ (ie, 6%-10%). Of note, our cohort included only 2 cases of primary testicular DLBCL, which most likely contributes to the lower prevalence of S1PR1+ cases in our study, compared to the previously reported studies mentioned above. The prevalence of S1PR1+ HGBL cases in our study appears to be similar to that reported in the literature for cases of DLBCL which are not primary testicular DLBCL.

In prior studies of pSTAT3 in DLBCL^16, 26, 27^, phospho-STAT3 (Tyr705) staining was found in 32-37% of DLBCL cases and appeared enriched in DLBCL, Non-GCB-type; one of these studies reported a higher prevalence (59%) of pSTAT3 positivity in DLBCL^16^. High nuclear expression of STAT3 and phosphoSTAT3 have been associated with unfavorable prognosis in DLBCL subgroups in these studies. Interestingly, S1PR1 has been reported to be transcribed by pSTAT3 and S1PR1 reportedly can, in turn, activate STAT3 in a positive feedback loop^15^. Furthermore, S1PR1 has been reported to be an effective target to block STAT3 signaling, tumor cell growth and metastatic spread using primary lymphoma cell models and DLBCL cell lines *in vitro* and *in vivo*^17^; however, the potential relationship between S1PR1 and pSTAT3 has not been adequately studied in clinical material from patients with DLBCL and other aggressive B cell lymphomas. In our cohort, we found that 38% of DLBCL cases (39/103) were positive for pSTAT3. This is in line with several prior studies mentioned above (32-37%). In addition, we found that HGBL also showed a similar prevalence of pSTAT3+ cases (35%, 9/26). Our study agrees with prior studies where pSTAT3 was expressed more frequently in Non-GCB-type, DLBCL cases compared to GCB-type, DLBCL cases^16, 17, 26, 27^. Lastly, our BL cases appeared to be uniformly negative for pSTAT3, and therefore, appear different from DLBCL and HGBL. Given the low number of S1PR1+ cases in our study, any potential relationship between S1PR1 and pSTAT3 expression could not be assessed in our cohort. Importantly, Koresawa et al^5^ did not observe a relationship between S1PR1 positivity and STAT3 phosphorylation among DLBCL cases by IHC analysis of FFPE sections. However, in a subset of their cases with frozen tissue available, Koresawa et al^5^ did observe a correlation between S1PR1 expression and phosphorylation of STAT3 by western blot analysis. Similarly, Liu et al^17^ also observed a correlation between S1PR1 expression and phospho-STAT3 using *primary* tumor cells from 10 Non-GCB-type, DLBCL patient samples and 2 DLBCL cell lines. These reported findings underscore the known difficulty of immunostaining of FFPE for phospho-epitope markers such as p-STAT3 due to tissue fixation and tissue processing factors^5^ and should be taken into consideration for future studies that explore the potential relationship between S1PR1 and pSTAT3 expression from clinical material.

FOXP1 is a transcription factor that typically functions as a transcriptional repressor and tends to be more highly expressed in DLBCL, Non-GCB-type (up to 71% of cases) compared to GCB-type cases^20, 22^. Moderate to high expression of FOXP1 in a high percentage of tumor cells has been associated with DLBCL, Non-GCB-type and a trend toward inferior outcome in DLBCL^20^. In terms of the S1PR2/FOXP1 axis, Flori et al^22^ examined FOXP1 expression and the relationship to S1PR2 expression in cell lines of both GCB-type and Non-GCB-type DLBCL. They found that strong expression of FOXP1 was more commonly observed in Non-GCB-type cell lines compared to GCB-type cell lines. Through FOXP1 siRNA knockdown in cell lines, followed by RNA sequencing and FOXP1-specific antibody chromatin immunoprecipitation, Flori et al identified S1PR2 as pro-apoptotic factor (ie, a tumor suppressor) both *in vitro* and *in vivo*, which is directly repressed by FOXP1. They also noted that the expression of S1PR2 was inversely proportional to FOXP1 expression in publicly available gene expression profiling studies from patient samples. Importantly, they found that high FOXP1 and low S1PR2 transcript levels were associated with inferior survival. The tumor suppressive properties of S1PR2 had also been previously reported by Muppidi et al^24^ who found that in mouse models, heterozygous loss of S1pr2 led to marked expansion of germinal centers, that could be repressed by overexpression of wild type S1PR2, but not a mutant S1PR2 incapable of signaling thru Gna13 and Arhgef1 to regulate cell growth and migration. They also found evidence for the importance of the S1PR2/GNA13/ARHGEF1 pathway in human DLBCL cell lines. This was consistent with earlier mouse model work by Green et al^23^ which also revealed an important role of S1pr2 in germinal center B cell homeostasis, whereby S1pr2 expression in germinal center B cells regulated apoptosis of germinal center B cells and also regulated germinal center B cell confinement via Gna12/Gna13/Arhgef1. Lastly, early studies by Cattoretti et al^6^ also found evidence that S1pr2 could act as a tumor suppressor in mouse models where S1pr2-deficient mice showed a tendency to develop lymphoma (GCB-type DLBCL) with aging. In terms of FOXP1 in our study, we have found that BL cases showed a unique pattern of expression with strong expression on FOXP1 in almost BL cases, while these BL cases lacked expression of S1PR1, pSTAT3, and S1PR2. The absence of pSTAT3 staining in Burkitt Lymphoma (a tumor of postulated germinal center B cell origin) appears consistent with the previously reported association of pSTAT3 expression in Non-GCB type, DLBCL. Also, the inverse expression pattern of FOXP1 and S1PR2 would appear to be consistent with the findings by Flori et al^22^ whereby expression of FOXP1 and S1PR2 were inversely proportional. The uniform expression of FOXP1 in our BL cases is somewhat interesting given that FOXP1 has been reported to be highly expressed in the context of Non-GCB type, DLBCL (although expression in GCB-type DLBCL has been described). Among our HGBL cases, we found that FOXP1 was expressed in 96% (25/26) of cases and in our DLBCL cases, we found that FOXP1 was expressed in 81% (83/102) and was not significantly different between GCB-type and Non-GCB-type DLBCL subgroups. The high prevalence of FOXP1 + HGBL cases (which is slightly higher than the prevalence of FOXP1+ DLBCL cases and approaches the prevalence of FOXP1+ BL cases in our cohort), is compatible with the concept that HGBL cases often demonstrate “Burkitt-like” cytogenetic (ie, MYC translocation), morphologic and immunohistochemical features, which are intermediate between DLBCL and BL.

Lastly, S1PR2 was expressed in approximately 19% (5/26) of HGBLs and in the case of Burkitt-like lymphoma with Del11q, but was negative in DLBCLs and BLs. The detectable expression of S1PR2 may at first appear counterintuitive, given that it is a putative tumor suppressor gene; however, loss of function mutations in S1PR2 have been described in DLBCL^24^, and for other biomarkers (eg, TP53), a precedent for detectable immunostaining in the presence of inactivating mutations has been described^28, 29^. Genotyping data of S1PR1 and S1PR2 are not available in our cases. Interestingly, in some of our case material, it was noted that S1PR2 was most strongly expressed in areas with high levels of apoptosis (eg, Figure 1: Germinal Center; Figure 2, panel K). Overall, among cases positive for S1PR staining, S1PR1 and S1PR2 staining were mutually exclusive in most cases (89%, 8/9), which is compatible with the prior concept that these receptors having antagonistic effects on cell growth and migration in several different cell systems^30^. As a category, HGBL comprised the majority of cases positive for S1PR1 or S1PR2 (78%, 7/9 cases). Given the low number of S1PR+ cases in our cohort, there was no clear relationship with MYC, BCL6 or BCL2 translocation status or with immunostaining for pSTAT3 or FOXP1. Eighty-nine percent (8/9) of S1PR+ cases were extranodal lymphomas, compatible with the reported findings in DLBCL where S1PR1 expression was associated with extranodal lymphomas^5, 16^.

**Figure 2:**
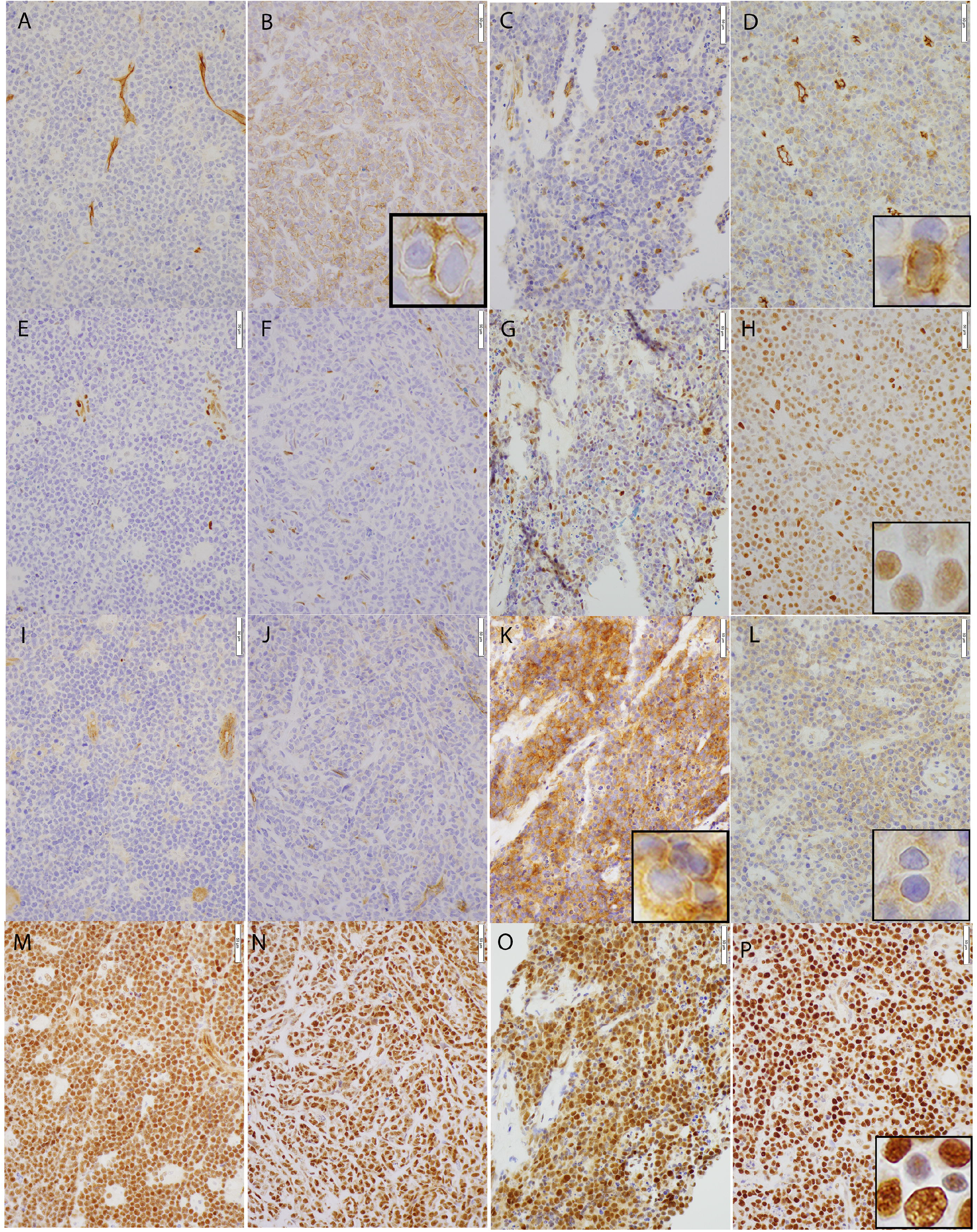
Representative immunostaining for S1PR1 (**A-D**), pSTAT3 (**E-H**), S1PR2 (**I-L**), and FOXP1 (**M-P**) in 4 different cases: Burkitt Lymphoma (**A, E, I, M**); HGBL-DH (**B, F, J, N**), HGBL,NOS (**C, G, K, O**) and HGBL,NOS (**D, H, L, P**). Burkitt lymphoma shows positivity for FOXP1. A HGBL-DH case (**B, F, J, N**) shows positivity for S1PR1 (**B**) and FOXP1 (**N**). A HGBL,NOS case (**C, G, K, O**) shows positivity for S1PR2 (**K**) and FOXP1 (**O**). A separate HGBL,NOS case (**D, H, L, P**) shows variable positivity for S1PR1 (**D**), pSTAT3 (**H**), S1PR2 (**L**) and FOXP1 (**P**). Images are shown at 40x magnification. Scale bars = 50um.

One of the main limitations of our study is the small size of the cohort which, given the low prevalence of S1PR1 and S1PR2 immunostaining, limits the interpretation of the findings. Additional studies are required on larger cohorts of these lymphoma types, in order to more fully explore the potential relationship between S1PR1/pSTAT3 and S1PR2/FOXP1 in clinical samples. In addition, studies with genotyping data available may also be helpful to helpclarify any potential relationships between mutation status (ie, genotype) and immunostaining (ie, phenotype) of S1PR1 and/or S1PR2.

In conclusion, herein, we report the staining patterns of S1PR1, pSTAT3, S1PR2 and FOXP1 in a cohort of aggressive, mature B cell lymphomas. We have found that: *i)* S1PR1 and S1PR2 showed different patterns of expression in mantle zones and follicle centers in reactive lymphoid tissue, *ii*) Burkitt lymphomas showed a unique pattern of expression compared to HGBL and DLBCL, *iii*) S1PR1 and S1PR2 were expressed in a low proportion of cases, which were predominantly HGBL involving extranodal sites and S1PR1, S1PR2 staining was predominantly mutually exclusive, *iv*) FOXP1 was expressed in a high proportion of the various case types and, *v*) pSTAT3, which represents a possible therapeutically targetable pathway, was detected in a significant proportion of HGBL and DLBCL cases. Taken together, these findings provide further evidence that S1PR1, pSTAT3, S1PR2 and FOXP1 play a role in a subset of aggressive mature B cell lymphomas and, therefore, deserve future study.

